# Differences in GenBank and RefSeq annotations may affect genomics data interpretation for *Pseudomonas putida* KT2440

**DOI:** 10.1101/2025.05.27.656432

**Authors:** Guilherme Marcelino Viana de Siqueira, Thomas Eng, Aindrila Mukhopadhyay, María-Eugenia Guazzaroni

## Abstract

Annotations of genomic features are cornerstone data that support routine workflows in conventional omics analyses in *Pseudomonas putida* KT2440 and other organisms. The GenBank and the RefSeq versions of the annotated KT2440 genome are two popular resources widely cited in the literature; yet, they originate from distinct prediction pipelines and possess potentially different biological information that is often overlooked. In this study, we systematically compared the features present in these resources and found that approximately 16% of the total of KT2440 ORFs show differences in their predicted genomic positions across GenBank and RefSeq, despite sharing equivalent locus tag codes. Furthermore, we show that these discrepancies can affect the results of high-throughput analyses by processing a collection of RNAseq expression datasets utilizing both annotations. Our findings provide a comprehensive overview of the current state of available resources for genomics research in *P. putida* KT2440 and highlight a rarely addressed yet widespread potential pitfall in the literature on this organism, with possible implications for other prokaryotes.

**Importance:** Genome annotation databases often rely on different statistical models for their function predictions and inherently carry biases propagated into studies using them. This work provides a quantitative assessment of two popular annotation resources for the model bacterium *P. putida* KT2440 and their influence on data interpretation. As large-scale omics datasets are commonly used to inform experimental decisions, our results aim to promote awareness of the caveats associated with these computational resources and foster reproducibility and transparency in *P. putida* research.

## Introduction

*Pseudomonas putida* KT2440 is a model saprophytic Pseudomonad strain that has been widely studied due to its innate metabolic versatility, including the ability to degrade a vast array of chemicals, ease of genetic manipulation, and its latent safety as a non-pathogenic microbe (1–3). The first version of the *P. putida* KT2440 reference genome (accession AE015451), published in the GenBank database by Nelson and collaborators in the early 2000s (4), greatly contributed to this popularity and rapidly accelerated research on this strain, enabling the construction of several metabolic models (5–9), and the leveraging of *P. putida*’s genetic traits for bioengineering efforts (1).

After the publication of the original genome sequence, the National Center for Biotechnology Information (NCBI) incorporated this assembly in its own reference sequence (RefSeq) database under the accession NC_002947. The RefSeq database is an NCBI project that offers a curated set of high-quality, non-redundant reference records, combining chromosomal, transcript and protein information, as well as structural and functional information and other metadata, for viruses, microbes, organelles, and eukaryotic organisms (10).

In 2021, RefSeq included over 200,000 bacterial and archaeal genomes (11). To ensure the quality of these annotations, it relies on a unified pipeline, the “Prokaryotic Genome Annotation Pipeline” (PGAP), which combines homology-based and *ab initio* methods for predicting functional elements directly from sequences (12). As a means to standardize nomenclature and reduce redundancies across the entire database, in 2015 the RefSeq team took on the massive effort of reannotating all of the prokaryotic genome assemblies deposited in the database using PGAP version 3 (13). This was done again in 2017 following the release of PGAP version 4.1 (13). Currently, the RefSeq team adopts a rolling schedule in which the oldest live assemblies are automatically re-annotated daily to benefit from updates incorporated to the pipeline (11).

Unlike RefSeq records, GenBank records are not revisited as often. In 2016, the GenBank record of the KT2440 genome annotation received its only major update to date (accession AE015451.2). This version is based on a resequencing of the original 2002 strain, with the employment of different computational tools (AMIGene and Prodigal), RNAseq expression data, and the manual curation of over 1,500 genes for the structural re-annotation of the genome (14). The new annotation includes over 300 newly predicted genes compared to the original record and lists 5,786 features, the vast majority of which (5,564) are protein-coding genes, identified by genomic loci codes with a “PP_” prefix. In contrast, the latest RefSeq annotation for *P. putida* at the timing of writing this manuscript (accession NC_002947.4), from December 2023, accounts for 5,693 annotated features, 5,486 of which are protein-coding genes that are named using a RefSeq “PP_RS” locus tag re-annotation prefix.

RefSeq is generally regarded as a reliable annotation source for microbial genomics (11), and the RefSeq annotation of the *P. putida* KT2440 genome is frequently cited in the literature. However, GenBank’s locus tag prefix format is often preferred to refer to specific genes in peer-reviewed publications — even when authors use recent RefSeq annotations — and the curated information provided by RefSeq appears to be unevenly propagated to other popular knowledge bases, including PseudomonasDB (15), BioCyc (16), Uniprot (17), KEGG (18), iModulons (19,20) and the Fitness Browser (21). Considering the way both annotation sources are used with the presumption they are truly interchangeable in the *P. putida* KT2440 literature, here we systematically compare the features present in each of them to assess the possible impacts of blindly choosing either for the interpretation of omics datasets in works utilizing *P. putida* KT2440 as a model strain.

## Materials and Methods

### *Cross-referencing* P. putida *KT2440 genome annotations*

*P. putida* KT2440 RefSeq and GenBank records were downloaded from the NCBI’s FTP server in September 2024 from assemblies GCF_000007565.2 and GCA_000007565.2, respectively. The total number of protein-coding genes in each annotation, as well as information on their genomic positions and other features of interest, were extracted from the \**_feature_table*.*txt*.*gz* file available in each directory. The resulting tables were processed in R (version 4.4.2) using the Tidyverse suite of packages (version 2.0.0) (22). The retrieval of data from KEGG to compose **Table 1** was made using the KEGGREST R package (version 1.44.1) (23).

**Table 1.** Overview of some of the main online genome resources for *P. putida* KT2440.

| Database | Associated annotations | Notes | Website |
| --- | --- | --- | --- |
| ALEdb | AE015451; NC_002947 | Includes other annotations of KT2440 genomes, depending on the experiment source. | https://aledb.org/ |
| BioCyc | NC_002947 | A 2016 version of the RefSeq annotation was used to populate the *P. putida* database, created in 2017. | https://biocyc.org/ |
| Fitness Browser | AE015451 | Provides their own set of curated re-annotations for over 40 *P. putida* genes based on new experimental data. | https://fit.genomics.lbl.gov/ |
| KEGG | AE015451 | Out of 5,786 entries (genes) on KEGG, only 1,804 are associated with metabolic pathways | https://www.kegg.jp/ |
| iModulonDB | AE015451 | Categorized the genome of *P. putida* KT2440 into 84 groups of independently modulated genes utilizing a machine-learning approach | https://imodulondb.org/ |
| UniProt | AE015451 ; NC_002947 | Offers hyperlinks to both GenBank and RefSeq for any given accession, although RefSeq locus tags themselves are not recognized as search terms. | https://www.uniprot.org/ |

The functional categorization of position-shifted genes was performed using annotations from the COG database (24). The treemap visualization for these functional annotations (**Figure 3**) was made using the package treemap (version 2.4.4) (25) in R. The overrepresentation analysis for COG annotations was conducted using a one-tailed Fisher’s exact test. All of the data processing steps and the code used in this analysis are available on Github (https://github.com/GuazzaroniLab/ppuGenomeAnnotations).

### Collecting historical data on genome annotations usage in the literature

Citation trends for the RefSeq and GenBank *P. putida* KT2440 genome annotations were obtained from Google Scholar search results using the terms “NC_002947” and “AE015451”, respectively. The metadata from all of the indexed publications was captured using a custom R script using RSelenium (version 1.7.9) (26). The resulting database was curated to remove duplicate entries, preprints, and other low-quality results. The final table is available as a plain text file (**Supplemental File 1**). The code used to retrieve and process publication metadata is available on GitHub (https://github.com/GuazzaroniLab/ppuLiteratureMetrics).

### Transcriptomic data processing

To compare the impacts of using either annotation source in the calculated results of a standard RNAseq pipeline, we downloaded data from several different publicly available RNAseq datasets from NCBI’s Sequence Read Archive (19,27–33) (**Supplemental Table S1**), composing a collection of different biological conditions and sequencing library layouts, and processed them using both the RefSeq and the GenBank versions of the transcriptome as reference.

In short, for this analysis, raw sequencing data (FASTQ format) were downloaded using the NCBI fasterq-dump command line utility (version 2.11.3). The retrieved data were then further processed for the removal of sequencing adapters and low-quality base assignments with Trimmomatic (version 0.39) (34). RSEM (35) was used to map reads to either the RefSeq or the GenBank version of the reference *P. putida* KT2440 transcriptome, and the DESeq2 package (version 1.44.0) (36) was used for determination of differentially expressed genes.

The code used for retrieving and processing the expression datasets for this analysis is available in GitHub (https://github.com/GuazzaroniLab/ppuRNAseqCollection). The resulting expression datasets are available as plain text files in **Supplemental File 2** (GenBank-processed) and **Supplemental File 3** (RefSeq-processed).

## Results

### Both GenBank and RefSeq genome annotations are widely used in the literature

To verify historical trends in the usage of the GenBank or RefSeq *P. putida* KT2440 genome accession codes in the literature, we retrieved Google Scholar results for the queries “AE015451” and “NC_002947”, respectively. As of December 2024, out of the roughly 400 selected publications retrieved after curation, 178 (43%) included the GenBank *P. putida* KT2440 genome accession code, whereas 238 (57%) publications included the RefSeq identifier. Historical data shows that RefSeq has always been more predominant than the GenBank in number of citations by a small margin (**Figure 1A**), but this seems to have become more prominent around 2016, coinciding with the release of the RefSeq annotation version 4 (NC_002947.4), the current major version as of the timing of the writing of this manuscript. In total, the RefSeq version of KT2440’s genome annotation has received four major updates since 2002, and a total of 65 version changes in total (**Figure 1B**). The GenBank record, as previously noted, is not updated as often, and has received only one major version change in 2016 (14), which is still the current version.

**Figure 1.**
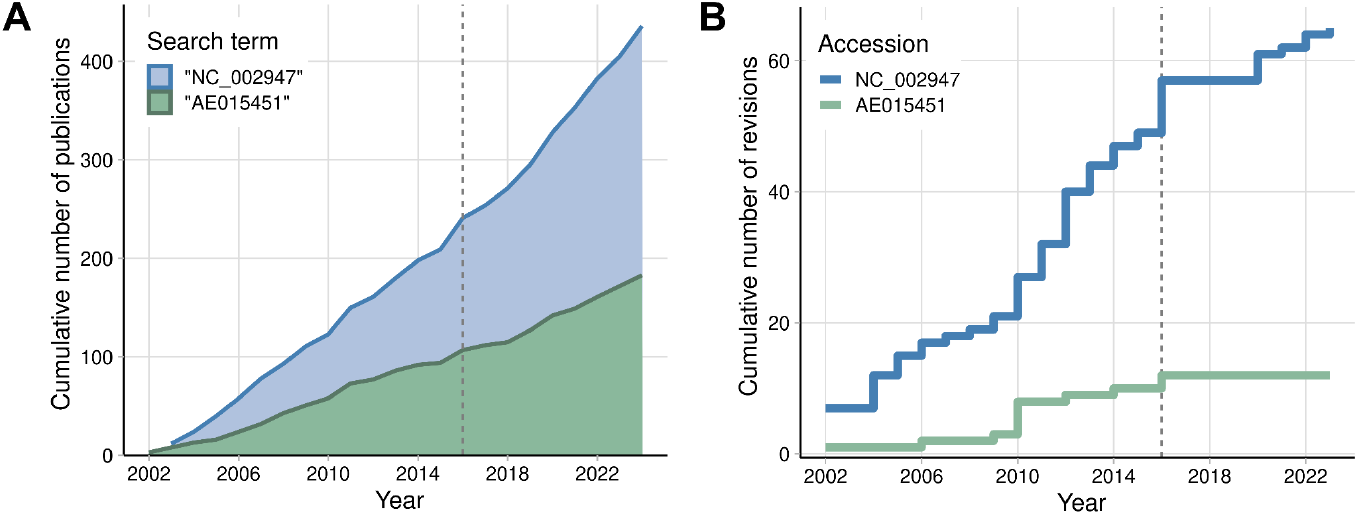
Historical trends for citations and the version history of GenBank and RefSeq *P. putida* KT2440 genome annotations. **A)** Google Scholar search results for publications containing the search terms “NC_002947” (RefSeq) or “AE015451” (GenBank) over time. **B)** Revision history of the RefSeq (blue) and GenBank (green) genome annotation records in the Nucleotide NCBI database (nuccore). In both panels, the year 2016, when both NC_002947.4 and AE015451.2 were introduced, is indicated by the vertical grey dashed line.

KT2440 genes are historically associated with a “PP” or “PP_” prefix followed by a four-digit number (4,14). Owing to the Prokaryotic RefSeq Genome Re-annotation initiative announced by NCBI in 2015, recent RefSeq annotations introduce a new locus tag code system for their own assigned ORFs using the prefix “PP_RS” followed by a five-digit number (37). Due to the many updates to the entire RefSeq database, and the difficulty in retrieving older versions from the website, it is hard to ascertain when the RefSeq version of the *P. putida* KT2440 genome annotation started to include this new locus tag system. However, the PGAP annotation pipeline provides a useful direct correspondence between new RefSeq locus tags and the GenBank annotation tags whenever an equivalent CDS is re-annotated on the new record. For instance, under this new nomenclature, PP_0001 is equivalent to PP_RS00005, PP_0002 to PP_RS00010, etc.

Relying on these conversions can however become a source of confusion as the gene products for a given locus tag might receive different nomenclatures across different knowledge bases depending on the underlying source it pulls data from (**Table 1**). For instance, several hypothetical proteins in GenBank were later reannotated with more specific nomenclature using PGAP, such as PP_1173 (*porin-like protein* in GenBank, and *OprD family porin* in RefSeq), and PP_2747 (*conserved protein of unknown function* in GenBank and *AMP-binding protein* in RefSeq). A search for these genes in UniProt will often match either the GenBank (in the case of PP_1173) or RefSeq (in the case of PP_2747) names, whereas in databases like KEGG they only coincide with the GenBank nomenclature. Moreover, the new RefSeq locus tag codes are also not widely recognized as search terms in any of the mentioned databases. For instance, data from ALEdb, a database that aggregates data from several adaptive laboratory evolution projects (38), show that the vast majority of unique mutations recorded in the database come from genomes analyzed using the RefSeq NC_002947.x family of annotations as a reference (**Supplemental Figure S1**), even though the genes themselves are only searchable in the database with the GenBank code. This favors the maintenance of the GenBank codes as the standard way to refer to specific loci in the *P. putida* KT2440 genome, despite the possibility of converting between old and new locus tag annotations obscuring more pressing differences between both sources, as explored in more detail in the next sections.

### The genomic coordinates of an expressive amount of genes are different in GenBank and RefSeq

Even though it is possible to map annotations between RefSeq and GenBank, we find that both the *number* of features annotated as “protein-coding” (genes) and their *positions* can vary greatly between both resources, as seen in **Figure 2A**. Importantly, while the vast majority (5,045) of annotated genes are present in both, 897 annotations with equivalent locus tag codes differ either in the start or end genomic positions. Almost half of these position differences comprise 27 base pairs or fewer, although a few cases reach a couple hundred base pairs **(Figure 2B, Supplemental File 4)**, for loci distributed all over the genome **(Figure 2C)**. We found that the re-annotation did not introduce any frameshifts to ORFs, except for the transposases in loci PP_2977 and PP_4420. Yet, as a result, 455 annotated genes had a different start codon position and 442 had a different stop codon position between both annotations, effectively changing the transcript size predicted by each of them, with RefSeq ORFs being smaller in the majority of the cases (**Supplemental Table S2**).

**Figure 2.**
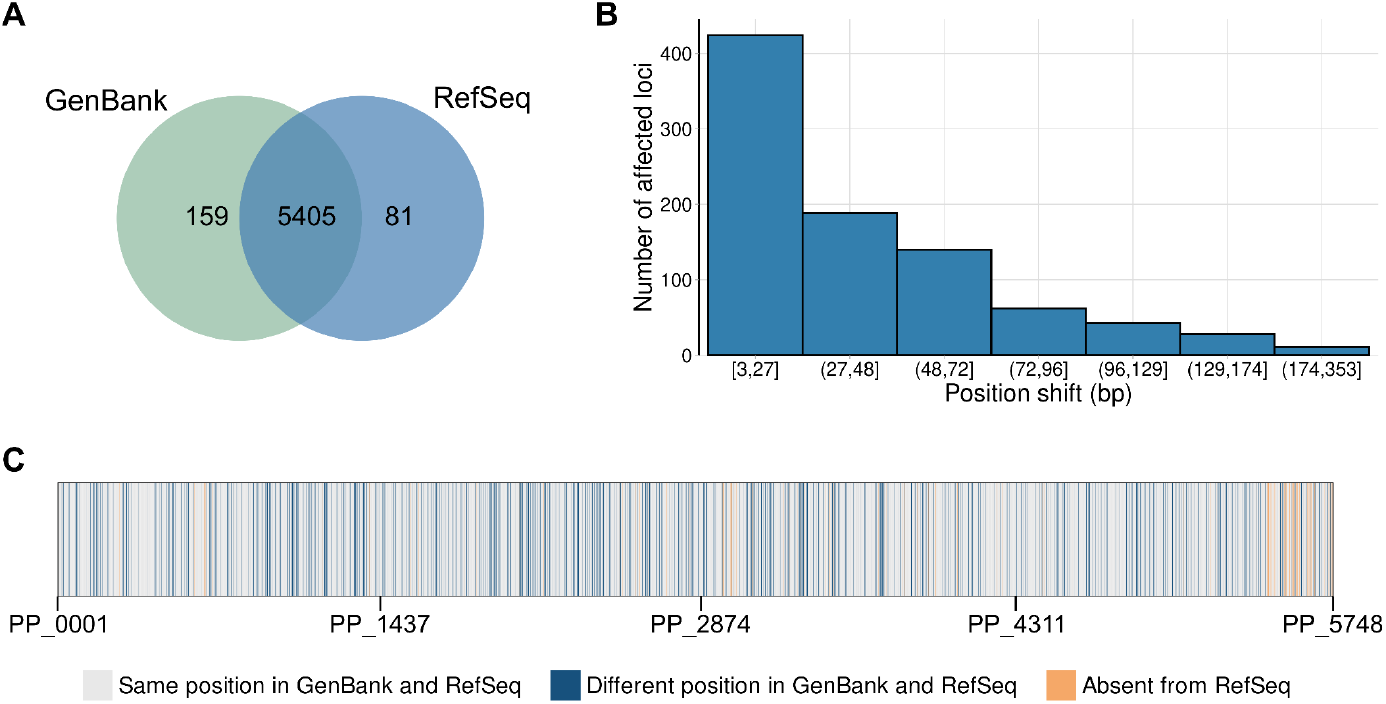
Comparison between features in GenBank and RefSeq *P. putida* KT2440 annotations. **A)** Venn diagram comparing the number of features (genes) in GenBank (green) and RefSeq (blue) with the “protein-coding” descriptor. **B)** Position differences in the start or end nucleotide of the shifted ORFs (in base pairs). **C)** Categorization of genes across all of the possible GenBank loci in comparison to their RefSeq counterpart. For these comparisons, RefSeq re-annotation locus tag codes were matched to the GenBank locus tag codes by using the “*old_locus_tag*” field whenever available. Please note that panel **C** does not accurately reflect the genome structure of *P. putida* KT2440, as locus tag codes do not necessarily match their relative genomic positions.

There are 159 genes in the GenBank annotation that were later excluded from the RefSeq annotation. On the other hand, RefSeq introduces its own set of 81 genes without a direct counterpart in GenBank. In both cases, most of the genes were annotated as conserved proteins of unknown function and hypothetical proteins. However, unlike Genbank or RefSeq exclusive genes, the set of genes with differences in their positions along the genome (or *shifted genes*) is much more diverse. It includes enzymes, transporters, and transcriptional regulators among other classes of proteins.

To better understand the diversity of biological functions associated with the shifted genes, we utilized the latest set of COG (Clusters of Orthologous Genes) classifications available in NCBI (24), and compared the classification of genes within the shifted set with those that remain in the same position in both annotations. Among the 897 shifted genes, only 260 did not have COG annotations associated with them (**Figure 3**). For those with associated annotations, we used Fisher’s exact test to verify whether any bias towards one of the five COG functional categories would be detectable in this subset. We found that genes belonging to the “cellular processes and signaling” category are overrepresented in the shifted subset in relation to the remainder of the genome (*p*-value < 0.001), with COG groups D (Cell cycle control, cell division, chromosome partitioning), U (Intracellular trafficking, secretion, and vesicular transport), Z (Cytoskeleton), N (Cell motility), V (Defense mechanisms) and O (Posttranslational modification, protein turnover, chaperones) more strongly associated with the shifted genes within this category (**Supplemental Figure S2**). As literature is heavily biased towards well-annotated genes, and given that annotation quality often relies on a small subset of experiments which cannot capture the full extent of protein functions (39,40), we believe that these results might inform neglected aspects of the *P. putida* KT2440 genome that could benefit from efforts to provide further context of their roles in this microbial host.

**Figure 3.**
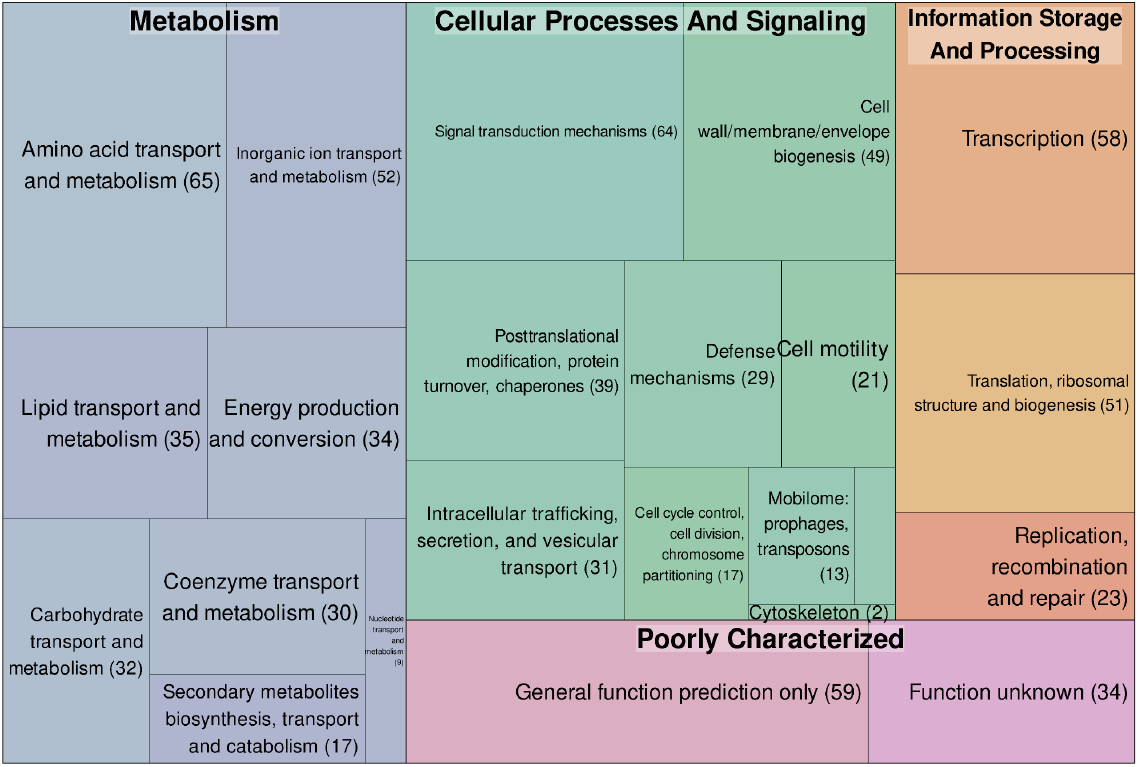
Cluster of orthologous genes (COG) classification for all functionally annotated shifted genes. The area of the squares correlates to the total number of genes associated with each category, which is also indicated in parenthesis. Smaller boxes with similar colors correspond to COG term descriptions that fit within the same functional categories (indicated in bold).

### The interpretation of expression data can be affected by the choice of using either annotation source

Considering that the length of the mRNA transcripts encoded by genes annotated differently between GenBank and RefSeq can vary, we sought to quantify the difference in choosing either reference annotation in the quantity and quality of differentially expressed genes found by a standard RNAseq pipeline (**Methods**). Under this analysis, a scatter plot of Log_2_ Fold Change values for the majority of the genes that are annotated in the same genomic coordinates in the GenBank and RefSeq forms a straight line along the center of the plot, suggesting that the calculated expression levels for genes of the same predicted length on both annotations tend to stay the same. Alternatively, genes with differently sized transcript sequences appear as a noisy cluster scattered across both axes, as illustrated in the four panels of **Figure 4**.

**Figure 4.**
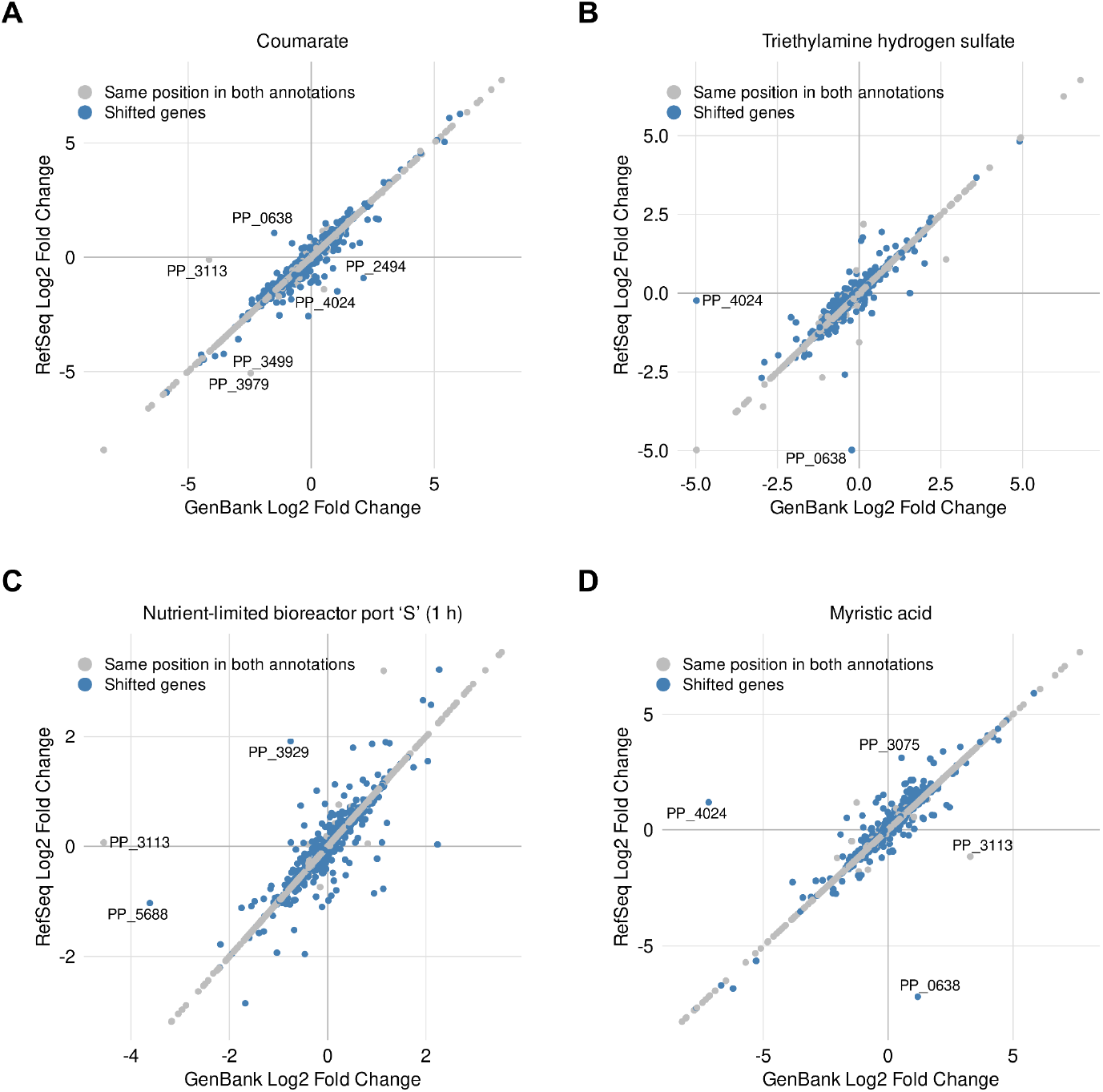
Comparison of calculated Log_2_ Fold Change values for shifted and unshifted *P. putida* KT2440 genes using RefSeq or Genbank annotations as a reference to calculate expression data. Panels **A–D** represent different experiment conditions retrieved from available RNAseq datasets (**Supplemental Table S1**). In each panel, blue circles represent shifted genes, whereas gray circles represent unshifted genes. Gene annotations using the conventional “PP_” prefix are given for all genes with an absolute difference greater than 2.5 between their RefSeq and GenBank Log_2_ Fold Change values. Among the highlighted genes, only PP_2494 (hypothetical protein), PP_3929 (hypothetical protein), PP_5688 ((2Fe-2S)-binding protein), and PP_3075 (sigma 54-interacting transcriptional regulator), all shifted genes, are not annotated as transposases.

Despite this variation, the differences in log_2_ fold change between both datasets were not too large even for shifted transcripts. In fact, the largest variations in log fold change typically belong to transposases (such as PP_3113, PP_3499, PP_3979, PP_4024, and PP_0638), regardless of whether transcript length changes between annotations. Proper transposon identification is a particularly challenging aspect of annotating both prokaryotic and eukaryotic genomes due to their repetitive nature and variable abundance (41–43), therefore it is unsurprising that this is one of the most prominent changes we found across sources that used different tools to generate gene annotations. Yet, due to the differences between both sources, the subset of genes considered to be significantly differentially expressed can differ between analyses, and includes diverse classes of genes (**Supplemental Figure S3**).

While it is beyond the scope of this work to benchmark different RNAseq analysis approaches, at minimum, we can conclude from this set of results that a gene can be classified as differentially expressed or not when either reference transcriptome is used to assign RNAseq reads to a transcriptome. We did not find examples of genes with altered significance status that were explicitly linked to validated phenotypes or discussed in the original publications we used as a source for this study. However, as publications rely more on the automatic processing of large amounts of data, these results reinforce that robust statistical analysis coupled with the use of sensible thresholds, ideally aided by experimental validation, remains a critical step for drawing meaningful biological conclusions from large datasets while avoiding spurious correlations inherent to them.

## Discussion

In 2019, Ghatak et al., introduced the term “y-ome” as the set of genes of an organism without experimental evidence of function, and described the y-ome of the textbook bacterium *Escherichia coli* K-12 MG1655 after curating data from several online resources (44). However, separating the “known” from the “unknown” relies on foundational assumptions about genes and genomic annotations that are often untested as researchers are often more interested in biological phenomena specific to their study. Here we found that not only the quality of functional annotations were uneven for *P. putida* KT2440 in available knowledge bases, but that the existence of two concurring major genomic annotations for this organism creates a divide in the literature (**Figure 1**), which, if unaccounted for, contributes to variable interpretation of data and confusion due imprecision in describing specific gene loci. For example, when validating a set of mutants for fatty acid and alcohol metabolism, Thompson and collaborators generated a *P. putida* KT2440 strain with a complete internal in-frame deletion of PP_2675, a gene that encodes for a type of cytochrome C protein involved in the assimilation of alcohols (45). In the GenBank version of the genome, the start codon of the downstream gene PP_2676 lies within the ORF of PP_2675, meaning that the resulting strain is an unintentional double mutant for both PP_2675 and PP_2676. On the other hand, in the RefSeq annotation the PP_2676 (PP_RS13935) start codon lies six base pairs downstream to the end of PP_2675 (PP_RS13930), and there is no overlap between the transcripts. We found this scenario to occur at over one hundred loci across our dataset (**Supplemental File 5**), and while the specific case of PP_2675 was acknowledged in a correction issued shortly after this first publication (46), determining which version of the ORFs corresponds to the biologically expressed *in vivo* remains challenging since they both derive from *in silico* predictions.

Such discrepancies require additional experimental efforts to fully characterize phenotypic behavior in *P. putida* under different experimental conditions, which, in turn, hinders progress in elucidating new biological functions in this host. Despite further published examples addressing these differences being hard to locate, 897 genes, over 16% of all of the annotated genes in *P. putida* KT2440, fall under the shifted category (**Figure 2**). As this subgroup is biased towards specific functional annotation categories (**Figure 3**), this effect may permeate research on KT2440. This is especially relevant considering that the broader context surrounding start codons can heavily influence protein expression in bacteria (47,48). If shifted genes are selected for heterologous expression, such as in metabolic engineering or synthetic biology approaches, these variations may lead to differing biological outcomes that are solely dependent on the dataset used, even if variables such as the Shine-Dalgarno sequence are controlled for. Moreover, while we have illustrated this problem for *P. putida*, it undoubtedly exists for all microbes where genome annotations are updated asynchronously from the research community.

Propagating and harmonizing annotations across databases is not a genomics- or *P. putida*-specific issue but rather a widely discussed challenge across multiple fields (49). Yet, these differences are often not fully appreciated by researchers working within the narrow scope of their model organism, even though the inadvertent use of different references may lead to varying interpretations of the same dataset (**Figure 4**). As household databases provide different versions of the *P. putida* genome annotation, obtaining a coherent picture of the interlinked knowledge of metabolic and physiological traits of this organism can be challenging. As a result, intense data-driven approaches that rely on information available from different sources, like large-scale metabolic or genomic models built by different groups, can produce conflicting results that require extensive manual curation to be reconciled (6,50). This effect is amplified when new taxonomic classifications for well-established strains or gene naming conventions, such as the RefSeq prokaryotic re-annotation project, or the recently proposed taxonomic re-classification of *P. putida* KT2440 as a *Pseudomonas alloputida* (51,52), break linkages of the known literature. All things considered, we hope the present work raises awareness of a latent but underappreciated problem in the *P. putida* genome annotation landscape, which should encourage researchers to deliberately choose, and faithfully describe, their specific reference materials in publications.

## Data availability statement

The source code used for analysis and figure generation in this work is available in GitHub repositories (see **Materials and Methods**). The BioProject accession codes for all RNA-seq datasets analyzed in this work are listed in the **Supplemental Table S1**.

## Acknowledgements

We thank our colleagues in the MetaGenLab and m-group research groups, as well as Tiago Cabral Borelli MSc., for their constructive comments on this work.

## CRediT Statement

Conceptualization: GMVS. Data acquisition, processing and interpretation: GMVS. Preparation of Figures: GMVS. Initial Draft of Manuscript: GMVS, TE. Editing of Manuscript: TE, GMVS, AM, M-EG. Raised Funds: M-EG, AM. All authors have read, given feedback, and approved the manuscript for publication.

## SUPPLEMENTAL INFORMATION

### Tables

**Table S1.** Summary information on the RNAseq datasets used in this work.

| BioProject Acc. | Base Culture Media | Growth Condition(s) | Publication<br>(Referenced in Main Text) | Reference Transcriptome in<br>Original Publication |
| --- | --- | --- | --- | --- |
| PRJNA480068 | minimal salt medium | Myristic acid | Not published | - |
| PRJNA796354 | minimal salt medium | Various carbon sources | (19) | AE015451.2 (GenBank) |
| PRJNA885112 | LB | Selenium | (27) | GCA_000014625.1<br>(*P. aeruginosa* GenBank acc.) |
| PRJNA630234 | minimal salt medium | Ionic liquids | (28) | NC_002947.4 (RefSeq) |
| PRJNA533248 | minimal salt medium | Nutrient-limited<br>bioreactor | (29) | NC_002947.4 (RefSeq) |
| PRJNA486005 | minimal salt medium | Various carbon sources | (30) | NC_002947.3 (RefSeq) |
| PRJNA341451 | minimal salt medium | Solvents (toluene) | (31) | NC_002947 (RefSeq) |
| PRJNA338603 | minimal salt medium | Osmotic and oxidative<br>stresses, antibiotics | (32) | NC_002947.3 (RefSeq) |
| PRJNA450701 | minimal salt medium | Zinc | (33) | PseudomonasDB<br>(Derived from GenBank) |

**Table S2.** Comparison of differences in ORF sizes for shifted genes between GenBank and RefSeq annotations.

| Number of affected loci | Affected extremity | Min difference (bp) | Max difference (bp) | Difference median (bp) | Outcome on relative ORF size |
| --- | --- | --- | --- | --- | --- |
| 75 | 3' | 3 | 114 | 31.5 | RefSeq is larger |
| 380 | 3' | 3 | 261 | 33 | GenBank is larger |
| 78 | 5' | 3 | 353 | 27 | RefSeq is larger |
| 364 | 5' | 3 | 297 | 30 | GenBank is larger |

### Figures

**Figure S1.**
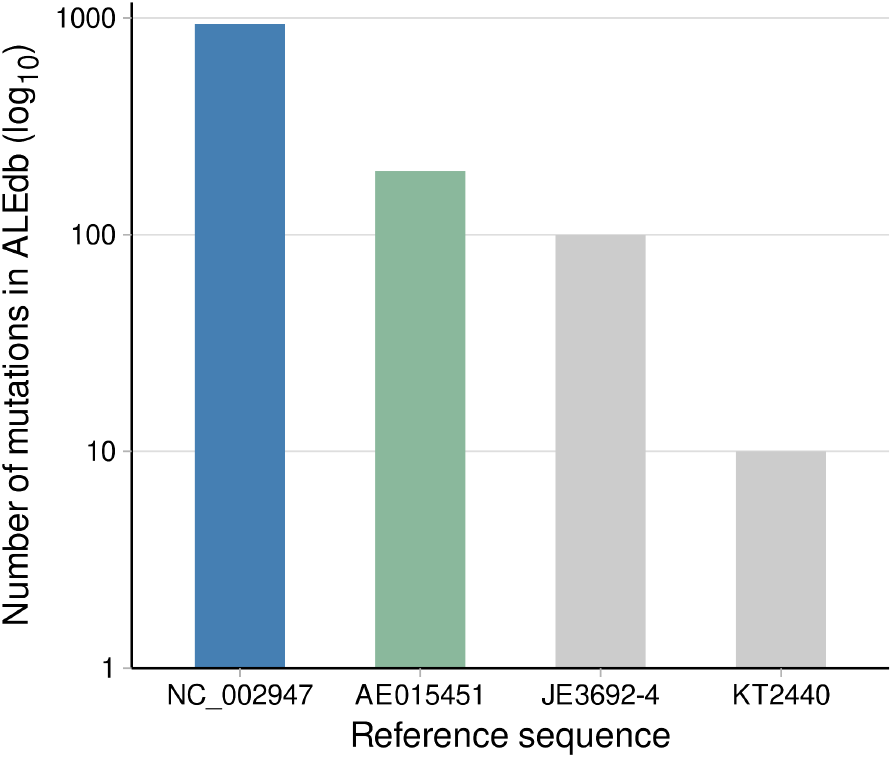
Reference genome annotation usage in ALEdb *P. putida* KT2440 mutation records. Data from ALEdb (https://aledb.org/) was downloaded using *P. putida*’s KT2440 taxonomic ID (160488) as the single search term in December 2024. Of note, there are 3 mutations in this database (at genomic positions 1,201,211; 3,702,766; and 4,993,315) that fall in the “grey zone” between annotations. This means that if the GenBank annotation of the genome is considered, the first two mutations are within ORFs (namely PP_1051, which encodes a Type II secretion system protein, and PP_3269, which encodes a general stress protein). Alternatively, if the RefSeq annotation is considered, only the third one is within the span of an ORF (PP_4402, which encodes the beta subunit of an alpha-ketoacid dehydrogenase).

**Figure S2.**
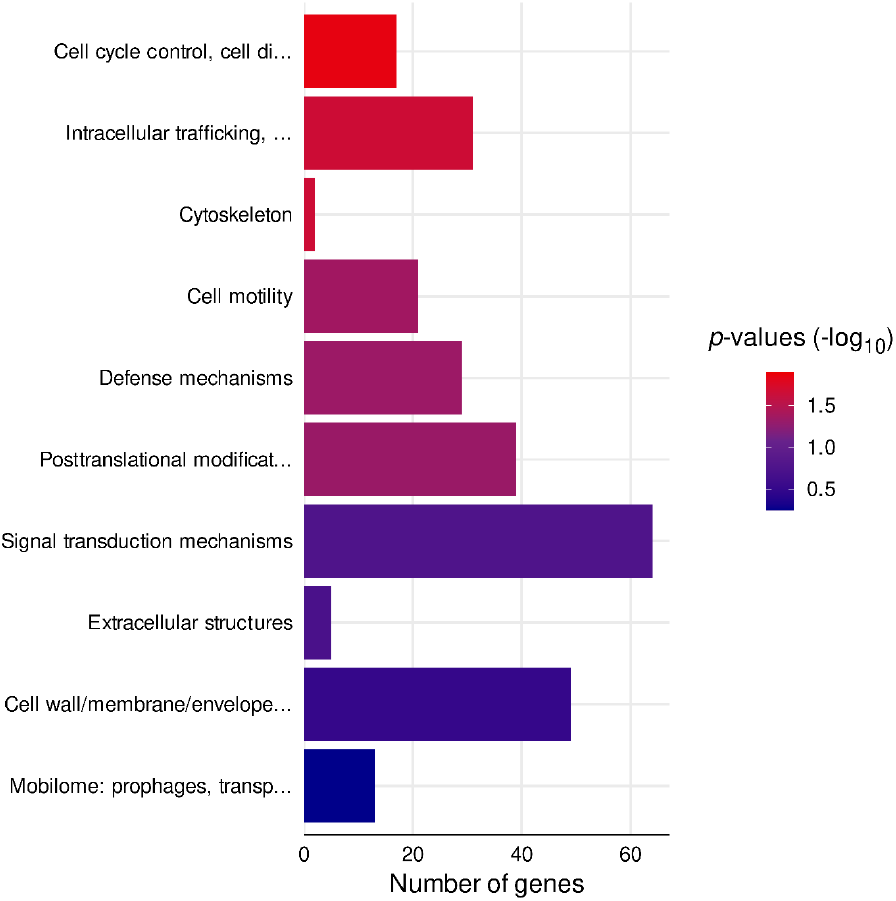
Overrepresentation analysis for the cluster of orthologous genes (COG) categories of shifted genes annotated with “Cellular processes and signaling” functions. In this figure, the number of shifted genes within each of the ten COG categories in this group is represented by horizontal bars. The color scale indicates *p*-values from one-tailed Fisher’s exact tests, which assess the association between shifted and unshifted genes for each specific annotation relative to the rest of the genes with COG annotations.

**Figure S3.**
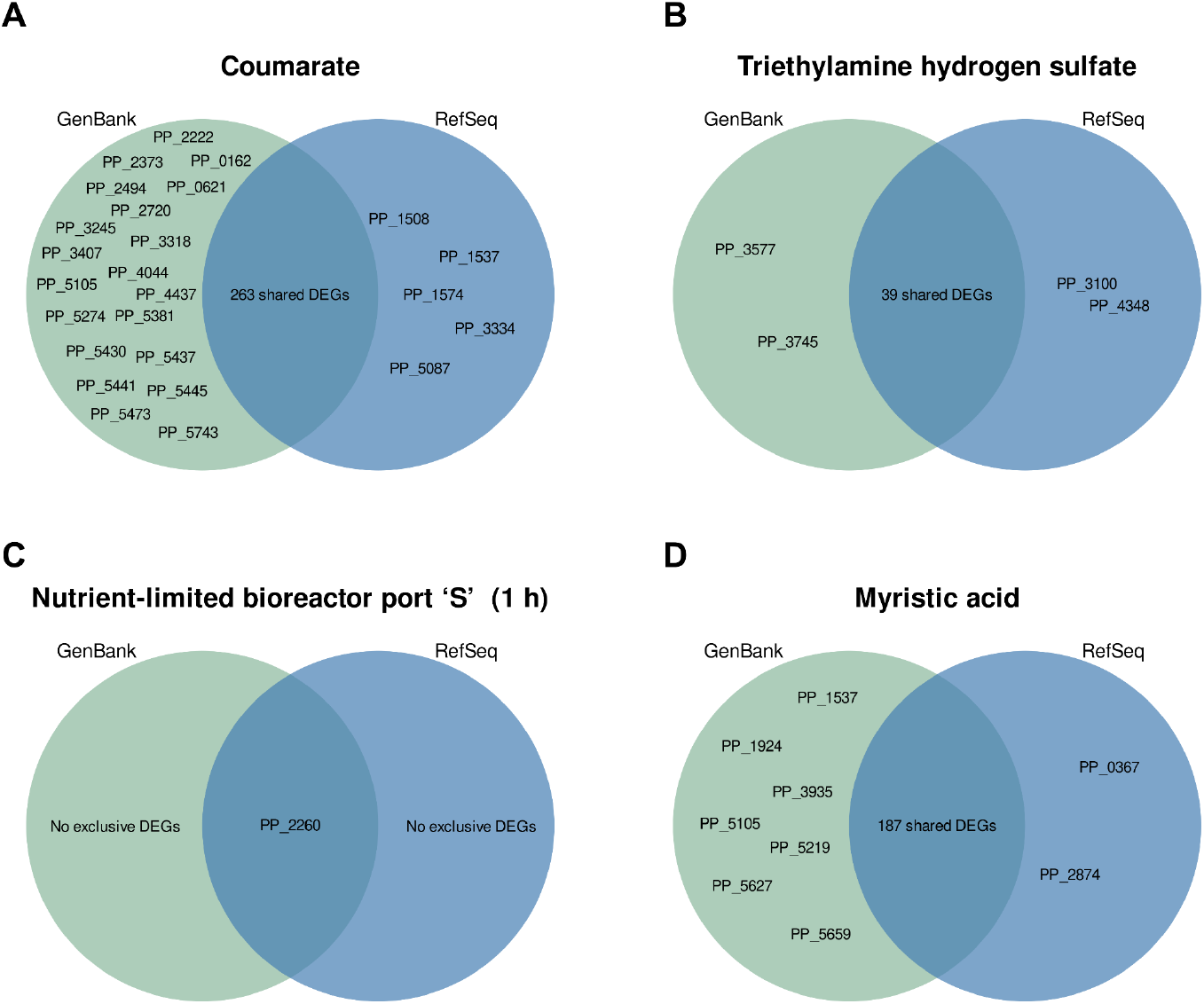
Differences in the output of RNAseq analyses due to the use of different reference transcriptomes. Venn diagrams in panels **A–D** compare the statistically significant differentially expressed genes (DEGs) detected in each RNAseq analysis using GenBank (green) or RefSeq (blue) as reference transcriptomes, considering adjusted *p*-values < 0.05 and absolute log_2_ fold change values ≥ 2 as thresholds. Due to the thresholds, the DEGs shown can either be up- or downregulated in the datasets. Data shown here derives from BioProject samples PRJNA796354 (19), PRJNA630234 (28), PRJNA533248 (29), and PRJNA480068 (unpublished) (**Table S1**). The panels in this figure match those of **Figure 4** in the main text.

